# β-H-Spectrin is a key component of an apical-medial hub of proteins during cell wedging in tube morphogenesis

**DOI:** 10.1101/2023.12.05.570290

**Authors:** Ghislain Gillard, Katja Röper

## Abstract

Coordinated cell shape changes are a major driver of tissue morphogenesis during development, with apical constriction or wedging of groups of epithelial cells for instance leading to tissue bending in folding or budding processes. During the budding of the tubes of the salivary glands in the *Drosophila* embryo we previously identified a key interplay between the apical-medial actomyosin that drives apical constriction with the underlying longitudinal microtubule array. At this microtubule-actomyosin interface a hub of proteins accumulates: in addition to the microtubule-actin crosslinker Shot and the minus-end-binder Patronin, we identified two actin-crosslinkers, β-H-Spectrin and Filamin, and the multi-PDZ protein Big bang as components of this apical-medial hub. Tissue-specific degradation of β-H-Spectrin led to reduction of apical-medial Big bang, F-actin, Shot and Patronin and concomittant defects in apical constriction and tube morphogenesis. Residual Patronin still present in the apical-medial position was sufficient to assist microtubule reorganisation into the longitudinal array. In contrast to Patronin and Shot, neither β-H-Spectrin nor Big bang required microtubules for their localisation. β-H-Spectrin instead appeared to be recruited to the apical-medial domain via binding to phosphoinositides that accumulated here. Overexpression of a β-H-Spectrin fragment containing its PH domain displaced endogenous β-H-Spectrin from the apical-medial domain and led to strong morphogenetic defects. The interconnected hub therefore required the synergy of membrane-associated β-H-Spectrin and microtubules and their respective interactors for its assembly and function in sustaining the apical constriction during tube invagination.

## Introduction

During development, tissues arise from simple precursors or primordia through complex shape changes. In many cases the primordia are simple polarised epithelial sheets that undergo in plane deformations such as convergence and extension as well as out of plane bending or budding deformations. At a cellular level, tissue bending can be induced by coordinated epithelial cell shape changes, going from a columnar or cuboidal shape to a wedge shape. An implementation of such cell wedging widespread throughout evolution occurs in the form of the apical constriction of epithelial cell, driven by apical actomyosin. Actomyosin accumulates within epithelial cells near cell-cell contacts, the adherens junctions, located at the apico-lateral side of cells, the so-called junctional pool. But especially during morphogenetic changes, epithelial cells also show a prominent apical-medial pool of actomyosin, just underneath the free apical surface, that is highly dynamic and pulsatile in its accumulation and has been shown to play a key role in the apical constriction observed in many tissues (Gillard & Roper, 2020; Martin *et al*, 2009; Munjal *et al*, 2015).

Apical-medial actomyosin undergoes cycles of actin network build-up and myosin recruitment, followed by contraction of the network and disassembly (Munjal *et al*., 2015). Because the actin network is connected to adherens junctions, each pulse of actomyosin contraction can pull on cell junctions and lead to a constriction of the cell’s apical surface, as long as a ratcheting mechanism or clutch stabilises the new smaller apical area. Otherwise the cycles of actomyosin contraction lead to apical area fluctuations but not shrinkage (Coravos & Martin, 2016; Martin *et al*., 2009). Pioneering work especially in *Drosophila* has identified several upstream regulators of this dynamic system, such as a GPCR ligand and several of its receptors that trigger a downstream cascade leading to relocation of a RhoGEF followed by Rho and Rho-kinase activation (Kerridge *et al*, 2016). This ultimately drives myosin phosphorylation and activation, but also affects actin regulators such as Diaphanous and hence the network the myosin works on (Mason *et al*, 2013).

We study the formation of the tubes of the salivary glands in the *Drosophila* embryo as a model process of tube budding from a flat epithelial primordium (Fig. 1A, A’ and Suppl.Fig. S1 A, B) (Girdler & Röper, 2014; Sidor & Röper, 2016). We previously identified two key cellular behaviours that in a highly coordinated spatio-temporal pattern drive the initiation of tissue bending as well as the continued invagination of all cells of the primordium (Sanchez-Corrales *et al*, 2021; Sanchez-Corrales *et al*, 2018). Cells near the invagination point undergo apical constriction driven by strong apical-medial actomyosin accumulation, whereas cells at a distance to the pit undergo convergence and extension in a radially arranged way to continuously feed more cells towards the invagination point. The directional neighbour exchanges in these cells are driven by the polarised accumulation of junctional actomyosin (Fig. 1A, A’). Interestingly, we showed that the apical-medial pool of actomyosin critically depends on the underlying microtubule cytoskeleton that in constricting cells becomes organised into a longitudinal array with minus ends anchored apically (Fig. 1B) by the microtubule-interacting proteins Shot and Patronin and the microtubule-severing protein Katanin (Booth *et al*, 2014; Gillard *et al*, 2021). If microtubules are depleted or the longitudinal array is not formed, as shown upon Patronin, Katanin or Shot depletion, then apical constriction fails to proceed as in the wild-type due to defects in the apical-medial recruitment of actomyosin (Booth *et al*., 2014; Gillard *et al*., 2021). Shot, the sole fly spectraplakin protein, is a huge protein with possibilities for many protein interactions, but in particular with an N-terminal actin-binding domain and a C-terminal microtubule binding domain, thus being able to crosslink the two cytoskeletal systems (Röper *et al*, 2002). Patronin and Shot have been shown to interact in other tissues as well as in vertebrate cells (Khanal *et al*, 2016; Nashchekin *et al*, 2016; Noordstra *et al*, 2016), and can further interact with the Katanin as well as Spectrin (Khanal *et al*., 2016).

**Figure 1.**
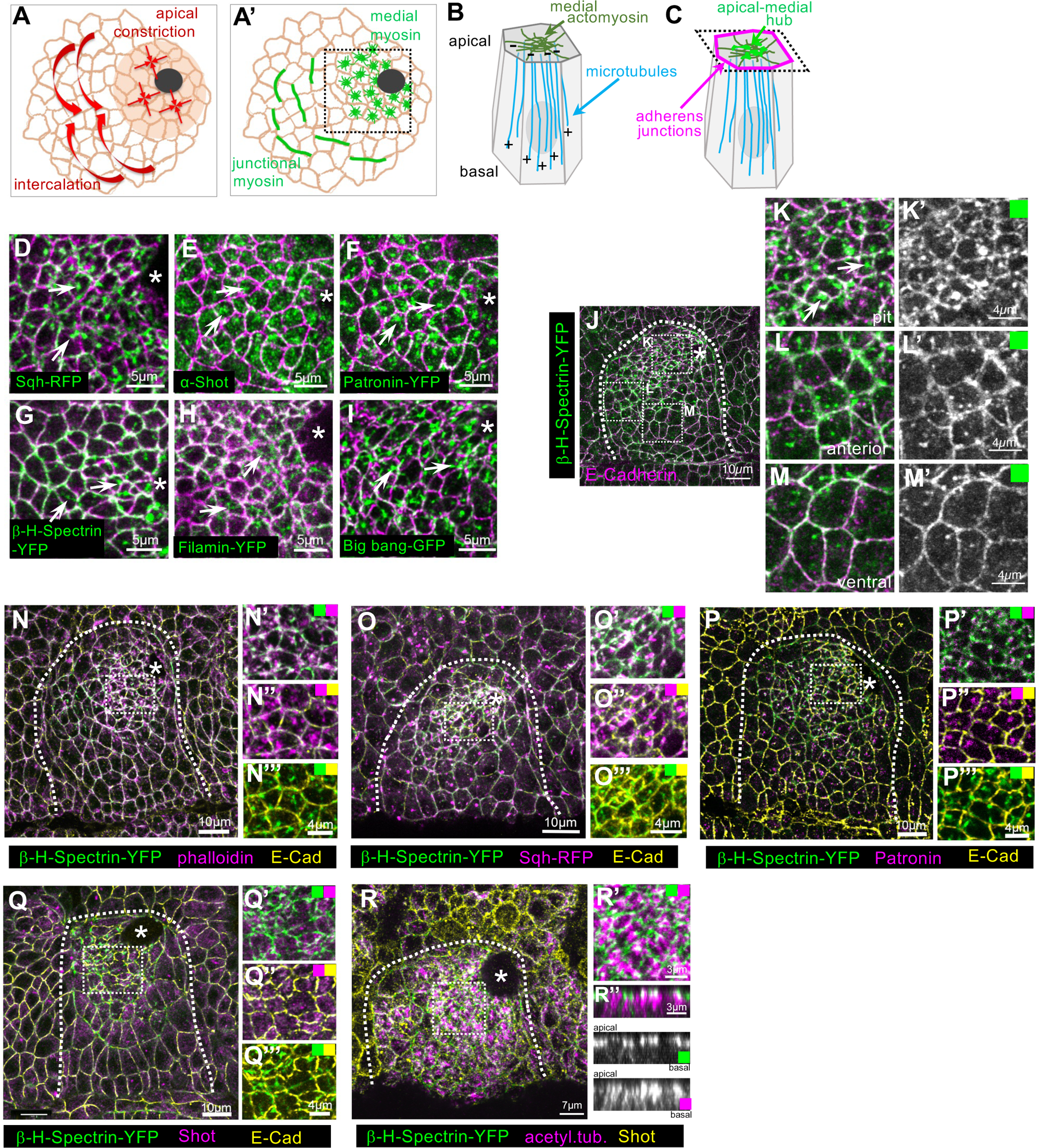
An apical-medial hub of proteins during apical constriction in tubulogenesis. **A, A’** Cells in the salivary gland placode during the initial steps of tubulogenesis show two distinct cell behaviours: apical constriction near the forming invagination pit (black circle) and directed intercalations (**A**). These are driven by distinct pools of apical myosin accumulation: apical-medial actomyosin in the apically constricting cells and polarised junctional accumulations in the cells undergoing intercalations (**A’**; (Sanchez-Corrales *et al*., 2021; Sanchez-Corrales *et al*., 2018)). **B, C** The apical-medial actomyosin is closely juxtaposed to the minus ends of a longitudinal non-centrosomal microtubule array in cells about to apically constrict, and this array and interaction between microtubules and actomyosin is crucial for constriction (Booth *et al*., 2014; Gillard *et al*., 2021). **D-I** Proteins localised to the apical-medial position in salivary gland placodal cells about to undergo apical constriction, forming an apical-medial hub (green): **D** myosin II (Sqh-RFP), **E** the spectraplakin Shot, **F** Patronin-YFP, **G** β-H-Spectrin-YFP (Karst-YFP), **H** Filamin-YFP (Cheerio-YFP), **I** Big bang-GFP. Apical membrane outlines are marked by E-Cadherin in magenta. **J-M’** β-H-Spectrin-YFP (green) is concentrated in the apical-medial position in cells close to the invagination pit, with a lower junctional contribution (**J**, **K, K’**), but is only found at the level of adherens junctions (marked by E-Cadherin, magenta) in non-constricting cells further anterior (**J**, **L**, **L’**) and further towards the ventral midline (**J**, **M**, **M’**). **N-R’’** Comparison of the apical-medial localisation of various components of the hub. **N-O’’’** β-H Spectrin-YFP (green) colocalises with phalloidin labelling F-actin (magenta; **N**-**N’’’**) and myosin (Sqh-RFP, magenta; **O**-**O’’’**) in the apical-medial position. **P-Q’’’** β-H Spectrin-YFP (green) also colocalises with Patronin (magenta, **P-P’’’**) and Shot (magenta, **Q-Q’’’**) in the apical-medial position. **R-R’’’** β-H Spectrin-YFP in the apical-medial position (green) colocalises with the ends of the longitudinal microtubule bundles marked by acetylated α-tubulin (magenta). Cell outlines are marked by Shot (yellow). Apical membranes in **N**-**Q’’’** are labelled by E-Cadherin (yellow), the white dotted boxes in **N**-**R** indicate the positions of the higher magnification two-colour images. Asterisks indicate the position of the invagination pit; dashed lines mark the boundary of the salivary gland placode. All immunofluorescence and live images of salivary gland placodes in this and subsequent figures are oriented with anterior side on the left and dorsal side up. See also Supplemental Figure S1.

α- and β-Spectrins in most epithelial cells form a submembraneous network, tied into the cortical actin cytoskeleton. Spectrins usually form heterotetramers of two α and two β subunits that can bind and crosslink membrane adaptors, transmembrane receptors as well as actin filaments directly (Liem, 2016). In epithelial cells, the distribution of α-Spectrin is homogeneous, whereas different β subunits are specific for the apical and basolateral domains. In *Drosophila* two β subunits exist, with β-Spectrin found basolaterally and β-heavy(H)-Spectrin restricted to the apical domain (Thomas & Kiehart, 1994). Typically, β-Spectrins in vertebrates associate with the adaptor protein ankyrin via a central region and with actin, α-Catenin and protein 4.1 via an N-terminal domain, and they also contain a C-terminal Pleckstrin homology (PH) domain that can mediate phospholipid binding (Leterrier & Pullarkat, 2022). β-H-Spectrin contains many more spectrin repeats than β-Spectrin but lacks the ankyrin-binding domain, instead an extended C-terminal domain (β-H-33) appears crucial for membrane association (Williams *et al*, 2004).

We identified that at the critical microtubule-actomyosin-membrane interface in epithelial cells undergoing apical constriction during tube budding a large number of proteins accumulate. In addition to Shot and Patronin, that we previously reported (Gillard *et al*., 2021), we found that the actin crosslinkers β-H-Spectrin/Karst and Filamin/Cheerio as well as the multi-PDZ adaptor protein Big Bang (Forest *et al*, 2018) also accumulated here. We identified β-H-Spectrin/Karst as a key component of this apical-medial hub of proteins and as required for apical-medial F-actin accumulation and apical constriction. In β-H-Spectrin’s absence Shot and Patronin were reduced in their localisation, but microtubules still became organised into a longitudinal array. However, this usual reorganisation of the microtubule network was not sufficient to trigger proper morphogenesis in embryos depleted for β-H-Spectrin, likely because the apical-medial accumulation of actomyosin and Bbg was also reduced. As opposed to other constituents of the apical-medial hub of proteins such as Shot and Patronin, the apical-medial accumulation of β-H-Spectrin as well as Bbg did not rely on MTs. Rather, overexpression of the C-terminal β-H-33 domain of β-H-Spectrin including the PH-domain, but not without it, lead to a specific loss of the apical-medial pool of β-H-Spectrin, and highly aberrant apical constriction and tube morphogenesis. This suggests that phospholipid-binding was key to the localisation of β-H-Spectrin to the apical-medial region. In summary, membrane-associated β-H-Spectrin and its interactors as well as microtubule-minus ends and interacting protein are required in synergy to support the apical-medial actomyosin in its function during cell constriction and tube morphogenesis.

## Results

### A dynamic apical-medial hub of components assembles during apical constriction-driven tube budding

During tube morphogenesis of the salivary glands in the *Drosophila* embryo, the cells that are about to internalise at any given point show a particular arrangement of their cytoskeletal systems: a dense network of apical-medial actomyosin in very close apposition to the minus ends of a longitudinal microtubule array that runs the apical-basal length of the cells (Fig. 1 B, C) (Booth *et al*., 2014; Gillard *et al*., 2021). In addition to myosin accumulation (Fig. 1 D), we previously found the spectraplakin Shot (Fig. 1E) and the microtubule minus end-binding protein Patronin (Fig. 1F) localised within the apical-medial region (Booth *et al*., 2014; Gillard *et al*., 2021), using endogenously-tagged versions of Patronin (Patronin-YFP; (Nashchekin *et al*., 2016) and the myosin regulatory light chain, Spagetti Squash (Sqh-RFP; (Ambrosini *et al*, 2019)) compared to antibody labelling for Shot (Fig. 1E; Suppl.Fig. 1C-H)(Röper & Brown, 2003). Using further protein trap lines tagging genes at endogenous loci, we now uncovered that two actin crosslinkers, β-H-Spectrin (Karst; Fig. 1G; (Thomas & Kiehart, 1994)) and Filamin A (Cheerio; Fig. 1H; (Sokol & Cooley, 1999)) also accumulated in an apical-medial position, as did the multi-PDZ domain protein Big bang that was previously reported to bind Spectrins (Fig. 1I; (Forest *et al*., 2018)). β-H-Spectrin, like the other components identified here, was strongly enriched at the apical-medial position in cells of the salivary gland placode that were actively constricting apically or about to do so, whereas other placodal and epidermal cells showed a junctional accumulation of β-H-Spectrin (Fig. 1J-M’). To assess whether all these components were in fact localised to the same apical-medial domain, we performed pair-wise comparisons, between β-H-Spectrin-YFP and the other apical-medial accumulating proteins. We first co-labelled β-H-Spectrin-YFP and the actomyosin cytoskeleton using phalloidin to label F-actin (Fig. 1N-N’’’), as well as myosin regulatory light chain (Sqh-RFP, Fig. 1O-O’’’). Similarly, β-H-Spectrin-YFP localisation was compared with microtubule minus ends, assessed through the endogenous localisation of Patronin (Fig. 1P-P’’’) and Shot (Fig. 1Q-Q’’’) or using an antibody against acetylated α-tubulin that accumulates at the apical end of placodal microtubules (Fig.1 R-R’’; (Booth *et al*., 2014)). In all cases in fixed samples the components colocalised within the same apical-medial foci. Thus, in epithelial cells of the salivary gland placode about to or already undergoing apical constriction, a hub of proteins assembles at the interface between the contractile apical-medial actomyosin and the minus-ends of the longitudinal microtubule array.

Apical-medial myosin driving apical constriction displays a very dynamic behaviour, undergoing pulsatile increases and decreases in intensity as well as flow behaviour underneath the free apical surface (Munjal *et al*., 2015), and this is for instance also mirrored by the behaviour of the apical actin network on which the myosin works (Dehapiot *et al*, 2020). To investigate whether the components of the apical-medial hub colocalise in time as well as in space, we collected time lapse movies of β-H-Spectrin-YFP in comparison to Sqh-RFP, Shot-EGFP and Patronin-RFP (Fig. 2A-D). In all cases, apical-medial β-H-Spectrin-YFP colocalised at all time points with apical myosin, Shot and Patronin (Fig. 2 B’, C’, D’). These data highlight the formation of a dynamic apical-medial hub that assembles and disassembles as cells undergo apical constriction in the salivary gland placode.

**Figure 2.**
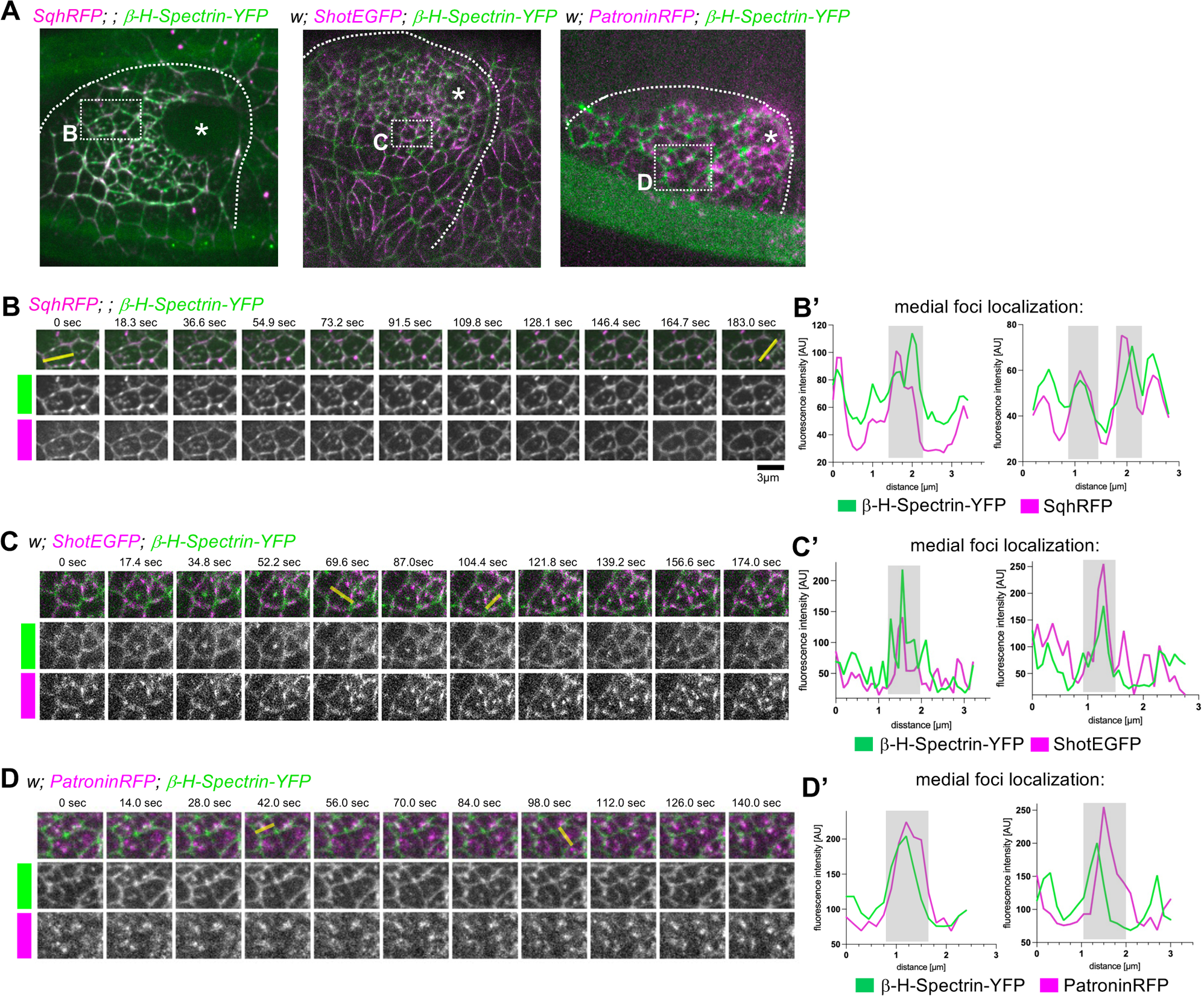
Proteins of the apical-medial hub show linked dynamics in the apical domain. **A** Still images of movies of two-colour time-lapse of *SqhRFP;; β-H-Spectrin-YFP*, of *w; ShotEGFP; β-H-Spectrin-YFP* and of *w; PatroninRFP; β-H-Spectrin-YFP* analysed further in higher magnification views in **B-D’**. Asterisks mark the position of the invagination pit, dotted lines mark the boundary of the placode and dotted boxes mark the position of the higher magnifications shown in **B-D**. **B** Close-up of individual placodal cells of *SqhRFP;; β-H-Spectrin-YFP* embryos, illustrating the colocalization of apical-medial foci of myosin (SqhRFP) and β-H-Spectrin-YFP, also quantified in line scans of individual cells in **B’**. **C** Close-up of individual placodal cells of *w; ShotEGFP; β-H-Spectrin-YFP* embryos, illustrating the colocalization of apical-medial foci of the two proteins, also quantified in line scans of individual cells in **C’**. **D** Close-up of individual placodal cells of *w; PatroninRFP; β-H-Spectrin-YFP* embryos, illustrating the colocalization of apical-medial foci of Patronin and β-H-Spectrin-YFP, also quantified in line scans of individual cells in **D’**. Grey boxes in **B’**, **C’**, **D’** indicate positions of apical-medial foci.

### β-H-Spectrin is required for apical constriction and the assembly and maintenance of the apical-medial hub

β-H-Spectrin-YFP showed a very distinct localisation to the apical-medial hub only in those cells about to or already undergoing apical constriction (Fig. 1 J-M’) and has been shown to be required for some aspects of apical constriction during gastrulation in *Drosophila* embryos (Krueger *et al*, 2020). We therefore set out to determine if β-H-Spectrin was required for this cell shape change within the salivary gland placode during tubulogenesis. In order to reduce β-H-Spectrin levels in a tissue-specific manner and not all throughout the embryo, we employed the UAS-degradFP system that targets YFP-or GFP-tagged proteins for degradation by the proteasome (Caussinus *et al*, 2012; Gillard *et al*., 2021) in combination with *fkhGal4* which drives specific expression in the salivary gland placode (Henderson & Andrew, 2000; Zhou *et al*, 2001). In control embryos (Fig. 3 A-B’ and E; *β-H-Spectrin-YFP fkhGal4*) β-H-Spectrin-YFP displayed a junctional pool in all epithelial cells and a strong apical-medial accumulation in cells undergoing apical constriction near the forming invagination pit (Fig. 3B’). When β-H-Spectrin-YFP was targeted for degradation (Fig. 3 C-E; *β-H-Spectrin-YFP fkhGal4 x UAS-degradFP; β-H-Spectrin-YFP*) the overall level of β-H-Spectrin-YFP was reduced throughout the salivary gland placode where *fkhGal4* was expressed, and whereas some junctional labelling remained, the apical-medial pool of β-H-Spectrin-YFP was strongly reduced (Fig. 3D’, E). We then analysed the effect this degradation of β-H-Spectrin-YFP had on apical constriction. In comparison to control placodes where apically constricting cells were clustered near the invagination pit (Fig. 3F) β-H-Spectrin-YFP-depleted embryos displayed fewer cells with apically constricted apices (Fig.3 G-H’).

**Figure 3.**
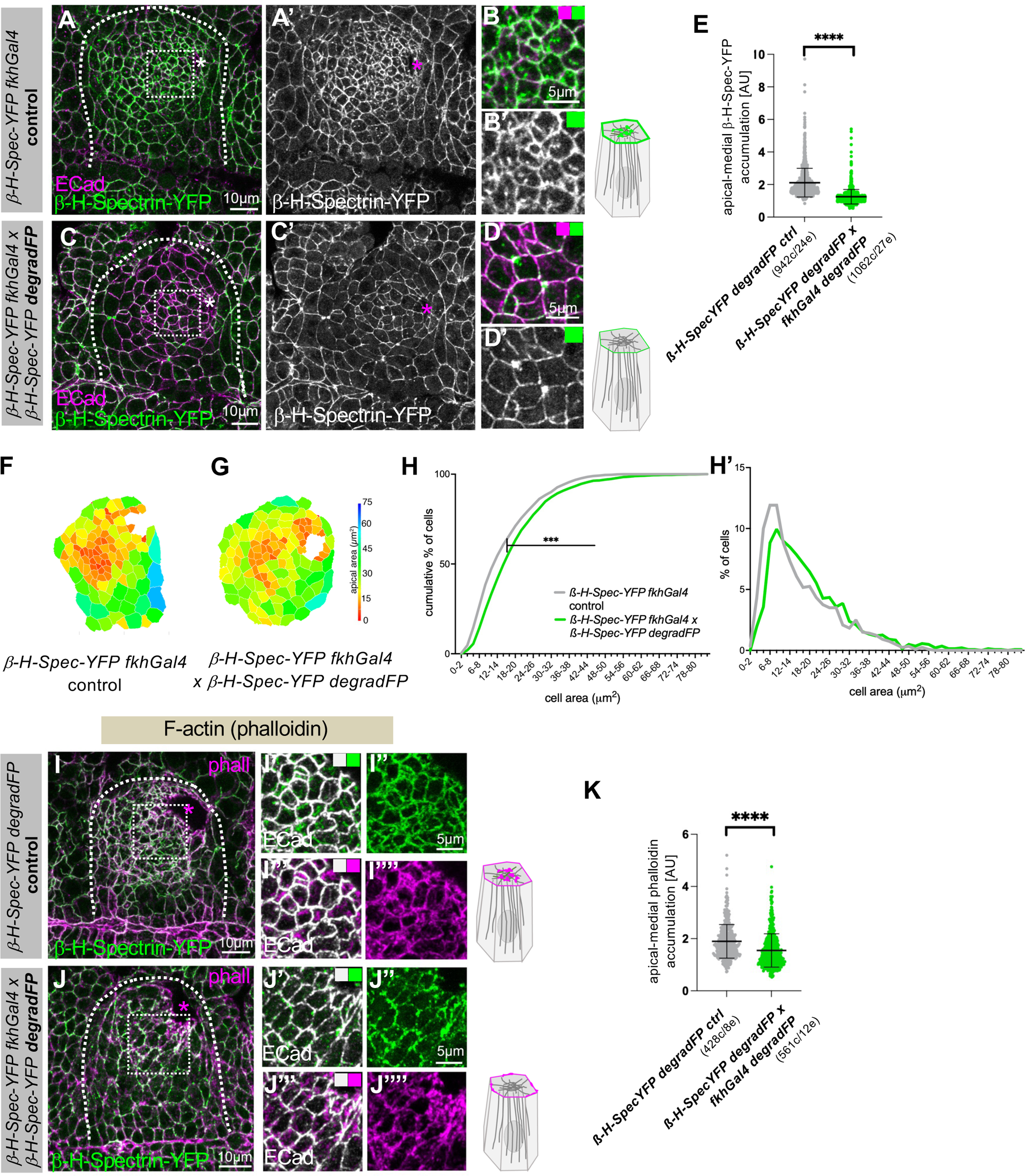
Loss of β-H-Spectrin impairs apical constriction. **A-E** In comparison to control placodes (*β-H-Spectrin-YFP fkhGal4*; **A-B’**) salivary gland tissue-specific degradation of endogenously tagged β-H-Spectrin-YFP (under control of *fkhGal4*) through the expression of an F-box/anti-GFP-nanobody fusion protein, degradFP (*β-H-Spectrin-YFP fkhGal4* x *β-H-Spectrin-YFP degradFP*) leads to significant loss of β-H-Spectrin-YFP (**C-D’**), in particular the apical-medial pool. Cell outlines are marked by E-Cadherin (E-Cad; magenta). **B, B’** and **D, D’** are higher magnifications of the white boxes marked in **A** and **C**, respectively. **E** Quantification of apical-medial β-H-Spectrin-YFP degradation (*β-H-Spectrin-YFP fkhGal4 control*; n=942 cells from 24 embryos; *β-H-Spectrin-YFP fkhGal4* x *β-H-Spectrin-YFP degradFP;* n=1062 cells from 27 embryos; shown are mean +/−SD, statistical significance was determined by two-sided unpaired Mann-Whitney test as p<0.0001). **F-H’** β-H-Spectrin-YFP degradation (**G**) leads to impairment of apical constriction compared to control (**F**), apical area of cells of example placodes are shown in a heat maps. **H, H’** Quantification of apical area distribution of placodal cells in control (*β-H-Spectrin-YFP fkhGal4 control*) and β-H-Spectrin depleted (*β-H-Spectrin-YFP fkhGal4* x *β-H-Spectrin-YFP degradFP*) placodes showing the cumulative percentage of cells relative to apical area size (**H**) and the percentage of cells in different size-bins (**H’**) [Kolmogorov-Smirnov two-sample test, P << 0.001 (***)]. 12 placodes were segmented and analysed for control and 15 for β-H-Spectrin-YFP depletion, the total number of cells traced was 1319 for control embryos and 1135 for *β-H-Spectrin-depleted embryos*. **I-K** β-H-Spectrin-YFP degradation (**J-J’’’’**) leads to reduction in apical-medial F-actin (phalloidin, magenta) accumulation compared to control (**I-I’’’’**). β-H-Spectrin-YFP is in green and E-Cadherin to label apical cell outlines is in white. **I’-I’’’’** and **J’-J’’’** are higher magnifications of the white boxes marked in **I** and **J**, respectively. **K** Quantification of apical-medial phalloiding in placodal cells in control (*β-H-Spectrin-YFP fkhGal4 control;* 428 cells from 8 embryos) and β-H-Spectrin depleted (*β-H-Spectrin-YFP fkhGal4* x *β-H-Spectrin-YFP degradFP;* 561 cells from 12 embryos) placodes. Shown are mean +/−SD, statistical significance was determined by two-sided unpaired Mann-Whitney test as p<0.0001. Asterisks indicate the position of the invagination pit; dashed lines mark the boundary of the salivary gland placode, and schematics indicate changes to the anlaysed component.

As β-H-Spectrin is an actin-binder and colocalised with apical-medial actomyosin, we analysed F-actin localisation in placodes where β-H-Spectrin was degraded (Fig. 3 I-K). In contrast to control placodes where F-actin displayed a strong medial pool (Fig. 3 I-I’’’), when β-H-Spectrin was degraded medial F-actin was also lost and more accumulated at junctions (Fig. 3 J-K). We also analysed the effect of β-H-Spectrin depletion on other components of the apical-medial hub of proteins. Previous work in a different tissue context had shown that β-H-Spectrin could also interact with both Shot and Patronin, two proteins we previously demonstrated localise to the apical-medial minus ends of microtubules and are key to proper apical constriction in the salivary gland placode (Booth *et al*., 2014; Gillard *et al*., 2021). β-H-Spectrin also interacts and collaborates with Bbg in wing discs (Forest *et al*., 2018). We therefore investigated the effect of β-H-Spectrin depletion on Patronin, Shot and Bbg localisation in placodal cells. In comparison to strong apical-medial foci of Patronin (visualised using endogenously tagged Patronin-RFP) in constricting salivary gland placodal cells of control embryos (Fig. 4A-A’’’’), when β-H-Spectrin-YFP was degraded we observed a reduction of apical-medial Patronin foci and an increased localisation at junctions (Fig.4 B-C). For both Shot and Bbg, the apical-medial foci observed under control conditions (Fig. 4D-D’’’’ and Suppl.Fig. S2A-A’’’) were also reduced when β-H-Spectrin-YFP was degraded (Fig. 4E-F and Suppl.Fig. S2B-C).

**Figure 4.**
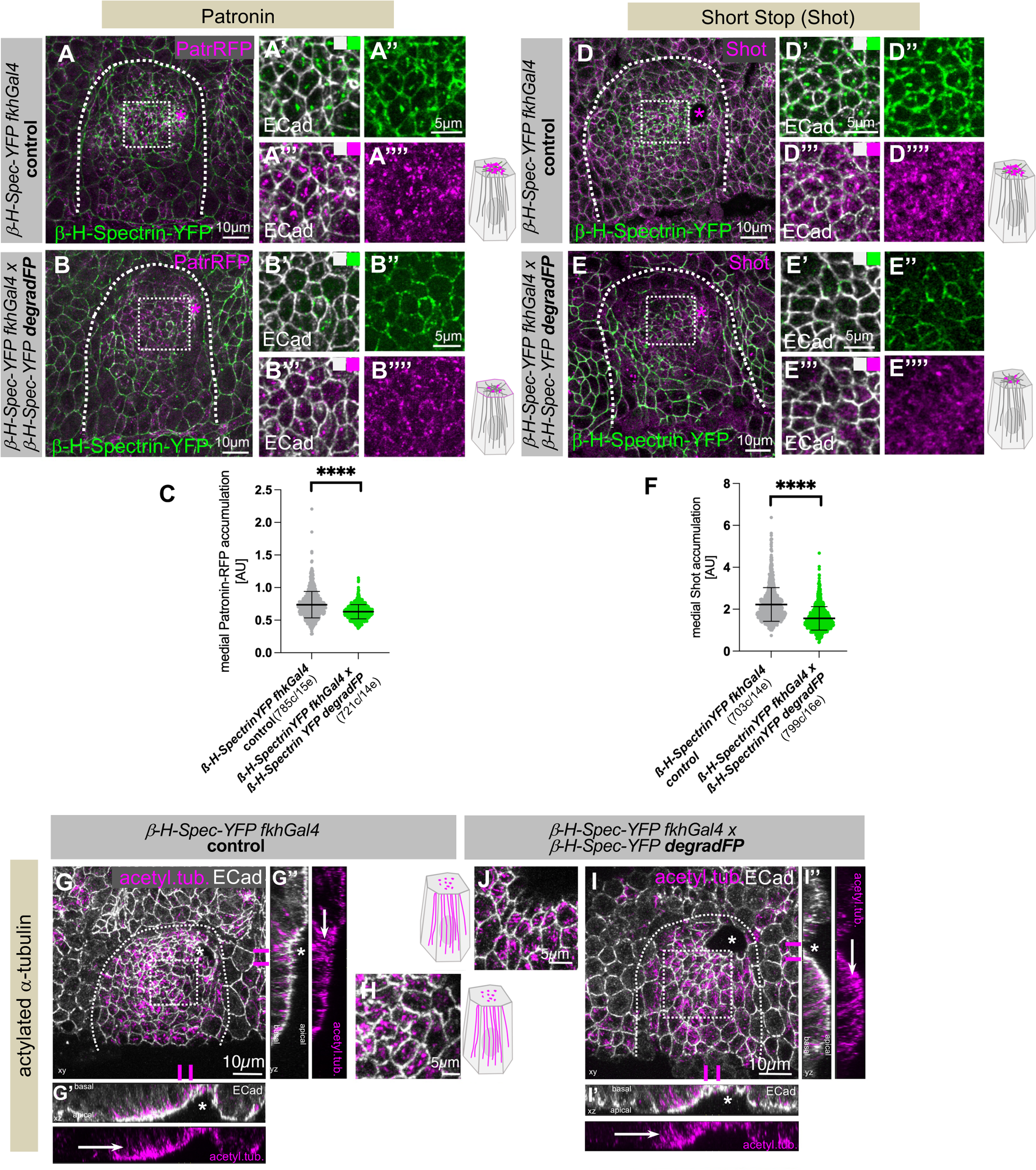
Loss of β-H-Spectrin leads to loss of the apical-medial hub. **A-C** β-H-Spectrin-YFP degradation (**B-B’’’’**) leads to a reduction in apical-medial Patronin foci (Patronin-RFP, magenta) compared to control (**A-A’’’’**). β-H-Spectrin-YFP is in green and E-Cadherin to label apical cell outlines is in white. **A’-A’’’’** and **B’-B’’’’** are higher magnifications of the white boxes marked in **A** and **B**, respectively. **C** Quantification of apical-medial Patronin in placodal cells in control (*β-H-Spectrin-YFP fkhGal4 control;* 785 cells from 15 embryos) and β-H-Spectrin depleted (*β-H-Spectrin-YFP fkhGal4* x *β-H-Spectrin-YFP degradFP;* 721 cells from 14 embryos) placodes. Shown are mean +/−SD, statistical significance was determined by two-sided unpaired Mann-Whitney test as p<0.0001. **D-F** β-H-Spectrin-YFP degradation (**E-E’’’’**) leads to reduction in apical-medial Shot (magenta) foci compared to control (**D-D’’’’**). β-H-Spectrin-YFP is in green and E-Cadherin to label apical cell outlines is in white. **D’-D’’’’** and **E’-E’’’’** are higher magnifications of the white boxes marked in **D** and **E**, respectively. **F** Quantification of apical-medial Shot in placodal cells in control (*β-H-Spectrin-YFP fkhGal4 control;* 703 cells from 14 embryos) and β-H-Spectrin depleted (*β-H-Spectrin-YFP fkhGal4* x *β-H-Spectrin-YFP degradFP;*799 cells from 16 embryos) placodes. Shown are mean +/−SD, statistical significance was determined by two-sided unpaired Mann-Whitney test as p<0.0001. **G-J** β-H-Spectrin-YFP degradation (**I-J**) does not affect formation of longitudinal microtubules (acetylated α-tubulin, magenta) that look comparable to control (**G-H**). **G’, G’’** and **I’, I’’** are cross section views at the positions indicated by magenta lines across the invagination pit. Asterisks mark the position of the pit, white arrows point to longitudinal microtubule bundles. **H** and **J** show higher magnifications of the white boxes marked in **G** and **I**, respectively, illustrating microtubule bundle ends visible within the apical domain as illustrated in the schematics. E-Cadherin to label apical cell outlines is in white Asterisks indicate the position of the invagination pit; dashed lines mark the boundary of the salivary gland placode. See also Supplemental Figure S2.

Thus, loss of β-H-Spectrin lead to concomitant changes in the localisation of other components of the apical-medial hub of proteins. We found previously that Patronin itself was recruited to the apical-medial region of placodal cells by microtubule minus ends, and that when Patronin was lost from these cells, the microtubule cytoskeleton did not form the longitudinal array. We therefore analysed the effect of loss of β-H-Spectrin on the microtubule cytoskeleton. Labeling for acetylated α-tubulin showed that in both control embryos (Fig. 4G-H and Suppl.Fig. S2D) as well as in embryos with β-H-Spectrin depletion in the placode (Fig. 4I-J and Suppl.Fig. S2E) microtubules in cells near the invagination point were organised into a longitudinal array with apical foci of microtubule bundles (Fig. 4 H, J) and extended bundles visible in cross-sections of placodal cells (Fig. 4 G’, G’’, I’, I’’). This suggested that the reduced levels of Patronin we observed in the absence of β-H-Spectrin did not prevent Patronin to perform its role in the reorganisation of the microtubule array. The levels of apical-medial Patronin were still sufficient to capture microtubule-minus ends released by Katanin, thereby generating the non-centrosomal microtubules that will then form the longitudinal array as we showed previously (Gillard *et al*., 2021).

These data illustrate that not only the longitudinal microtubule array but also apical-medial β-H-Spectrin are both required to support apical-medial actomyosin assembly and function to drive apical constriction.

### β-H-Spectrin is recruited to the apical-medal hub independently of microtubules

We now turned our attention to how β-H-Spectrin itself was recruited to the apical-medial position. Both Patronin and Shot do not only depend on β-H-Spectrin-YFP for their wild-type levels of apical-medial localisation but also depend on the presence of the longitudinal microtubule array in the constricting cells of the salivary gland placode (Booth *et al*., 2014; Gillard *et al*., 2021). The formation of the longitudinal microtubule array is in turn crucial to maintain the apical-medial pool of actomyosin (Booth *et al*., 2014). Therefore, in order to assess whether microtubules were required for β-H-Spectrin localisation we depleted microtubules in the salivary gland placode by overexpressing the microtubule-severing protein Spastin (Fig. 5; expressing *UAS-Spastin* under *fkhGal4* control). In both the control embryos (Fig. 5 A-A’’ and C; *β-H-Spectrin-YFP fkhGal4*) and when microtubules were depleted (Fig. 5 B-C; *β-H-Spectrin-YFP fkhGal4 x UAS-Spastin*) β-H-Spectrin-YFP was localised to apical-medial positions in apically constricting cells with no significant difference in the amount that accumulated.

**Figure 5.**
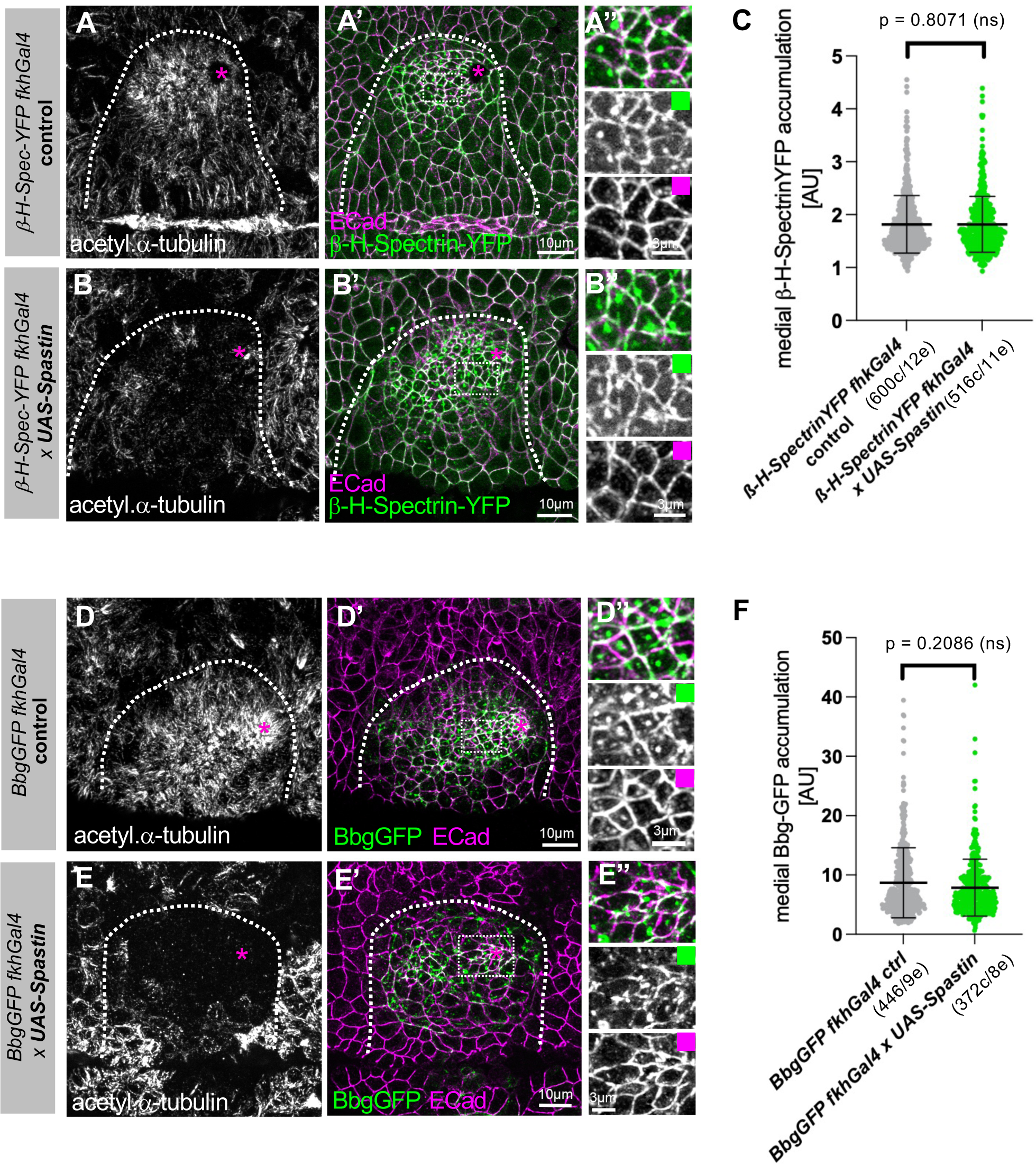
Microtubules are not required to recruit β-H-Spectrin or Big bang to the apical-medial hub. **A-C** In contrast to Patronin and Shot, localisation of β-H-Spectrin-YFP to apical-medial foci is not affected when microtubules are lost upon expression of *UAS-Spastin* under *fkh-Gal4* control (**A-A’’** for *β-H-Spectrin-YFP fkhGal4 control* and **B-B’’** for *β-H-Spectrin-YFP fkhGal4 UAS-Spastin*). **A’’** and **B’’** are higher magnifications of the white boxes marked in **A’** and **B’**, respectively. **C** Quantification of apical-medial β-H-Spectrin-YFP accumulation upon placodal microtubule loss (*β-H-Spectrin-YFP fkhGal4 ctrl:* 600 cells from 12 embryos; *β-H-Spectrin-YFP fkhGal4 x UAS-Spastin:* 516 cells from 11 embryos/ shown are mean +/− SD, statistical significance was determined by two-sided unpaired Mann-Whitney test as p=0.8071, non significant). **D-E’’** Localisation of BbgGFP to apical-medial foci is not affected when microtubules are lost upon expression of *UAS-Spastin* under *fkh-Gal4* control (**A-A’** for *BbgGFP fkhGal4 control* and **B-B’** for (*BbgGFP fkhGal4 UAS-Spastin*). **D’’** and **E’’** are higher magnifications of the white boxes marked in **D’** and **E’**, respectively. **C** Quantification of apical-medial BbgGFP accumulation upon placodal microtubule loss (*BbgGFP fkhGal4 ctrl:* 446 cells from 9 embryos; *BbgGFP fkhGal4 x UAS-Spastin:* 372 cells from 8 embryos/ shown are mean +/− SD, statistical significance was determined by two-sided unpaired Mann-Whitney test as p=0.2086, non significant). Microtubules are labelled in **A**, **B**, **D** and **E** with an antibody against acetylated α-tubulin that accumulates in the longitudinal array in placodal cells, E-Cadherin to label cell outlines is in magenta. Asterisks indicate the position of the invagination pit; dashed lines mark the boundary of the salivary gland placode.

Bbg had been shown by yeast-two hybrid and co-immunoprecipitation analysis to bind to β-H-Spectrin, an interaction that was necessary for Bbg’s stable localisation to adherens junctions in wing discs (Forest *et al*., 2018). We identified above that β-H-Spectrin was also key to Bbg’s robust localisation to the apical-medial hub in salivary gland placodal cells (Suppl. Fig. S2). We now tested whether Bbg also depended on an intact microtubule cytoskeleton for its apical-medial localisation in placodal cells or remained apical-medial, bound by β-H-Spectrin, when microtubules were depleted. Like β-H-Spectrin, in both the control (Fig. 5 D-D’’ and F; *BbgGFP fkhGal4*) and when microtubules were depleted (Fig. 5 E-F; *BbgGFP fkhGal4 x UAS-Spastin*) BbgGFP was localised to apical-medial positions in apically constricting cells with no significant difference in the amount that accumulated.

Thus, it appears that the apical-medial hub has two groups of components that either, like Patronin and Shot, depend on the underlying longitudinal microtubule cytoskeleton for their localisation or that, like β-H-Spectrin and Bbg, are independent of it and therefore are recruited in a different uncharacterised fashion.

### Phosphoinositide-binding via its PH domain contributes to β-H-Spectrin’s recruitment to the apical-medal hub

What other mechanism could mediate β-H-Spectrin’s recruitment to the apical-medial position? β-H-Spectrin’s C-terminal domain beyond the spectrin repeats, the β-H-33 fragment, contains a PH domain that can interact with inositol-phospholipids (Fig. 6A; (Williams *et al*., 2004; Zhang *et al*, 1995). We therefore investigated where phosphoinositides were localised within salivary gland placodal epithelial cells. We employed two transgenic markers of phosphoinositides: Grp1-PH-GFP, reported to bind PI(3,4,5)P_3_ (Fig. 5B-B’’’; *tub84B::grp1-PH-GFP*; (Britton *et al*, 2002)) as well as UAS-PLC∂PH-EGFP reported to bind PI(4,5)P_2_ (Fig. 6 C-C’’’; *fkhGal4 x UAS-PLC∂PH-EGFP*; (Zelhof & Hardy, 2004)). Classically, PI(4,5)P_2_ has been proposed to be important for apical membrane identity, whereas PI(3,4,5)P_3_ appeared enriched basolaterally in 3D cyst culture of MDCK cells (Martin-Belmonte *et al*, 2007; Roman-Fernandez *et al*, 2018). Recent studies during different processes of tissue morphogenesis in *Drosophila* have shown, though, that apical PI(3,4,5)P_3_ can also play an important role (Claret *et al*, 2014; Miao *et al*, 2021; Pinal *et al*, 2006). Ubiquitous expression of Grp1-PH-GFP revealed apical-medial pools of PI(3,4,5)P_3_ in constricting cells in the salivary gland placode, colocalising here with F-actin labelled by phalloidin (Fig. 6 B-C’’). Similarly, placode-specific expression of PLC∂PH-EGFP showed labelling of the apical-medial region and hence presence of PI(4,5)P_2_, in part overlapping with F-actin labelling by phalloidin (Fig. 6C-C’’’’). Thus, phosphoinositides were concentrated in a position that could support β-H-Spectrin recruitment.

**Figure 6.**
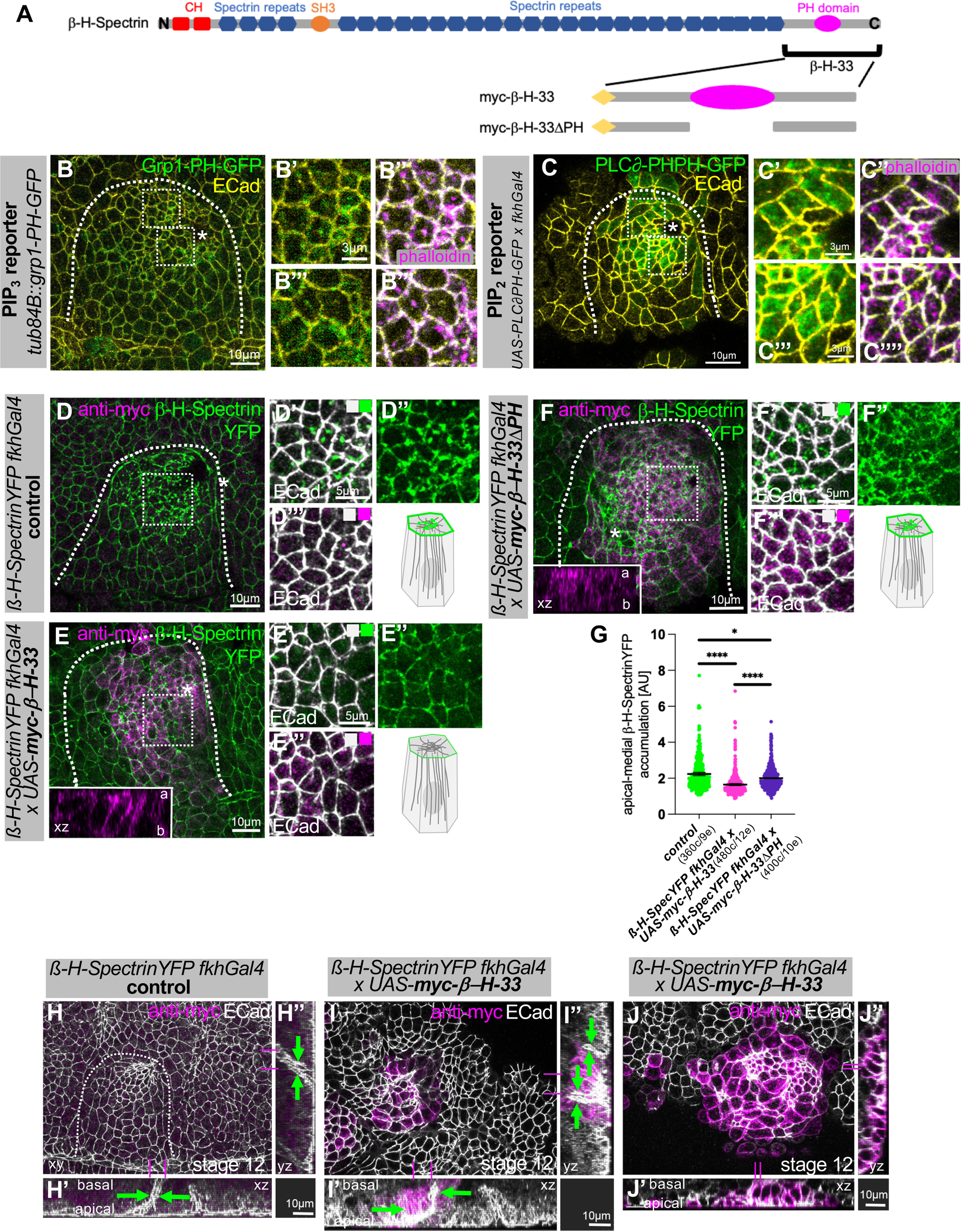
β-H-Spectrin’s localization to and function in the apical-medial hub depends on its PH domain. **A** Schematic of β-H-Spectrin protein domains, comprising N-terminal CH domains, 30 Spectrin repeats, an SH3 domain and the β-H-33 C-terminal domain that contains a phosphoinositide-binding PH domain. Indicated are also the parts of the C-terminal domain contained within overexpression constructs called *UAS-myc-β-H-33* and *UAS-myc-β-H-33ΔPH*. **B-C’’’’** Analysis of PI(3,4,5)P_3_ and PI(4,5)P_2_ localisation within the salivary gland placode, using the tagged PH domain of Grp1 as a PI(3,4,5)P_3_-sensor (*tub84B::grp1-PH-GFP*) and the PH domain of PLC8 as a PI(4,5)P_2_-sensor (*UAS-PLC8PH-GFP x fkhGal4*). **B-B’’’** Grp1-PH-GFP is in green, phalloidin to label F-actin, including apical-medial pools, is in magenta and apical cell outlines marked by E-Cadherin in yellow. Areas marked by white boxes in **B** are magnified in **B’**, **B’’**. **C-C’’’** PLC8-PH-GFP is in green, phalloidin to label F-actin, including apical-medial pools, is in magenta and apical cell outlines marked by E-Cadherin in yellow. Areas marked by white boxes in **C** are magnified in **C’**, **C’’**. **D-G** Overexpression of the C-terminal region of β-H-Spectrin, β-H-33, including or lacking the PH domain in the salivary gland placode. Expression of *UAS-myc-β-H-33* (**E**-**E’’’**) but not *UAS-myc-β-H-33ΔPH* (**F-F’’’**) in comparison to *fkhGal4* control (**D-D’’’**) leads to loss of apical-medial β-H-Spectrin-YFP accumulation. Insets in **E** and **F** show xz-cross-sections of placodal cells (a= apical, b= basal) illustrating the localisation of myc-β-H-33 (**E**) and myc-β-H-33ΔPH (**F**), with myc-β-H-33 localised to free apical and lateral membranes and myc-β-H-33ΔPH mostly cytoplasmic with minimal membrane localisation. **G** Quantification of apical-medial β-H-Spectrin-YFP accumulation upon *UAS-myc-β-H-33* or *UAS-myc-β-H-33ΔPH* expression compared to control (*β-H-Spectrin fkhGal4 ctrl:* 360 cells from 9 embryos; *β-H-Spectrin fkhGal4 x UAS-myc-β-H-33:* 480 cells from 12 embryos; *β-H-Spectrin fkhGal4 x UAS-myc-β-H-33ΔPH:* 400 cells from 10 embryos/ shown are mean +/− SD, statistical significance was determined by two-sided unpaired Mann-Whitney test as p<0.01 [*] and p<0.0001 [***]). Areas marked by white boxes in **D**, **E** and **F** are magnified in **D’-D’’’’**, **E’-E’’’’**, **F’-F’’’’**. Asterisks indicate the position of the invagination pit; dashed lines mark the boundary of the salivary gland placode. Schematics show the changes to endogenous β-H-Spectrin *observed.* **H-J’’** Overexpression of *UAS-myc-β-H-33* using fkhGal4 at a slightly later stage of tube invagination (embryonic stage 12) shows many poorly constricted cells and only cells with remaining apical-medial β-H-Spectrin-YFP still constricting (**I-J’’**; Suppl. Fig. S2) in comparison to control (**H-H’’**). Cross sections in **I’-J’’** illustrate aberrant or delayed invagination of the tube and aberrant or multiple lumen shapes (green arrows) compared to the single narrow lumen in control (green arrows in **H’**, **H’’**). Magenta lines indicate the position of cross section views. Dotted like in **H** indicates the boundary of the placode that in **I** and **J** is marked by transgene expression under *fkhGal4* control and anti-myc labelling (magenta).

We then aimed to disrupt β-H-Spectrin’s potential recruitment via phosphoinositides by expressing a construct comprising the β-H-33 fragment (Williams *et al*., 2004). Ectopic expression of this construct had previously been shown to interfere with normal salivary gland development (Williams *et al*., 2004), but the underlying reason was not clear. Expression of *UAS-myc-β-H-33* in the salivary gland placode under *fkhGal4* control, in comparison to control (Fig. 6D-D’’’ and G), led to a reduction of the apical-medial pool of β-H-Spectrin (Fig. 6E-E’’’ and G). myc-β-H-33 itself localised to the placodal cells’ plasma membrane (see cross section in inset in Fig. 6E) including the free apical domain (Fig. 6E’’’). By contrast, expression of a version of β-H-33 lacking the PH domain, *UAS-myc-β-H-33ΔPH*, under *fkhGal4* control did not affect the apical-medial pool of β-H-Spectrin (Fig. 6 F-F’’’ and G), and this version displayed a much more cytoplasmic distribution (see cross section in inset in Fig. 6F). Where the *myc-β-H-33ΔPH* construct localised to the plasma membrane apically it did not colocalise with β-H-Spectrin-YFP (Fig. 6F-F’’’). In slightly older salivary gland placodes (stage 12 rather than stage 11 as shown before), cells expressing high levels of myc-β-H-33 showed dilated apices, and only the few cells that retained apical-medial β-H-Spectrin foci appeared contracted (Fig. 6I-J’’), in contrast to the normal pattern of graded constriction observed in the control (Fig. 6H-H’’ and Suppl. Fig. S3 A-C’). Where portions of the placode managed to invaginate in myc-β-H-33 expressing placodes they formed an irregular-shaped tube (Fig. 6 I’,I’’).

These data suggest that β-H-Spectrin is, at least in part, recruited to the apical-medial region of placodal cells via its C-terminal PH domain, through interactions with phosphoinositides. We show that perturbation of this interaction through expression of a dominant-negative interactor, in the form of β-H-33, leads to aberrant apical constriction and aberrant tubulogenesis.

## Discussion

The submembraneous spectrin cytoskeleton has been studied in many different contexts since spectrins were originally identified as core components of the red blood cell cytoskeleton. In many instances, including in red blood cells but also, as more recently identified, in axons a key role for spectrins is membrane or cell stabilisation. This role is possible due to the expandable and hence reversibly stretchable structure of spectrin repeats. In both red blood cells and axons, the spectrin heterotetramers together with associated proteins such as membrane receptors and actin filaments are arranged in highly regular assemblies (Leterrier & Pullarkat, 2022; Teliska & Rasband, 2021). In epithelial cells by contrast, two β-Spectrin subunits are expressed and show a polarised distribution, with β-H-Spectrin confined to the apical domain and β-Spectrin localised basolaterally, as well as a less regular arrangement. Epithelial spectrins have been shown to play roles in epithelial morphogenesis in different organisms including in embryo elongation in *C.elegans* (Jia *et al*, 2019; McKeown *et al*, 1998), eye and follicle cell morphogenesis in *Drosophila* (Lee *et al*, 2010; Ng *et al*, 2016) and more recently also mesoderm invagination (Krueger *et al*., 2020). This requirement in dynamic developmental processes suggests that spectrins also serve important functions beyond stabilising and buffering a fixed cell shape.

We provide evidence for a very dynamic function of β-H-Spectrin in the rapid apical constriction of cells about to invaginate to form a tube, during the morphogenesis of the salivary glands in the fly embryo. Such apical constriction is driven by a highly dynamic and pulsatile pool of actomyosin within the apical-medial domain of cells, and β-H-Spectrin not only colocalises with but also dynamically co-pulsates with myosin and other components of this apical-medial hub. A related function has previously been proposed for β-H-Spectrin in mesoderm invagination in the fly embryo, though is not required for apical constriction *per se* but rather for apical ratcheting that allows the stabilisation of the cell apical domain after a pulse of actomyosin driven constriction (Krueger *et al*., 2020). We identify that β-H-Spectrin colocalises not only with actin and myosin at the apical-medial site, but also with the microtubule minus end-binding protein Patronin and the cytolinker Shot, as well as with the Filamin Cheerio and the scaffold protein Big Bang, thus forming an apical-medial hub of interacting proteins at the interface of actomyosin and microtubules (Fig. 7). We show that βH-Spectrin depletion reduces the apical-medial accumulation of Patronin and Shot. All three proteins have in fact been shown to directly interact in *Drosophila* follicle cells, with β-H-Spectrin acting as an upstream factor to recruit both Shot and Patronin at the apical membrane (Khanal *et al*., 2016). Shot and Patronin are required in the salivary gland placode to promote the formation of non-centrosomal microtubules, the anchoring of their minus-ends at the apical membrane and their orientation along the apical-basal axis. This non-centrosomal microtubule network is involved in actomyosin accumulation at the apical-medial site, a prerequisite for efficient apical constriction and tissue invagination (Booth *et al*., 2014; Gillard *et al*., 2021). The combined function of the hub of proteins at the apical-medial site is as yet unclear as all individually appear important for apical constriction, but βH-Spectrin and Filamin can crosslink actin filaments, Shot can crosslink actin and microtubules, and the other components can bind either actin or microtubules and also other components of the hub, generating a possibly highly interlinked structure (Fig. 7). Whilst this could suggest a more static function, the fact that all components appear highly dynamic and follow actomyosin’s pulsatile behaviour, the hub’s presence might allow recruitment of further factors with roles in the regulation of the dynamicity of apical-medial actomyosin. Furthermore, recent modelling approaches revealed that for a contractile and pulsatile actomyosin network a certain amount of crosslinking of actin is key (Belmonte *et al*, 2017). Therefore, components of the hub could directly influence the pulsations and contraction of the apical-medial actomyosin in the placodal cells.

**Figure 7.**
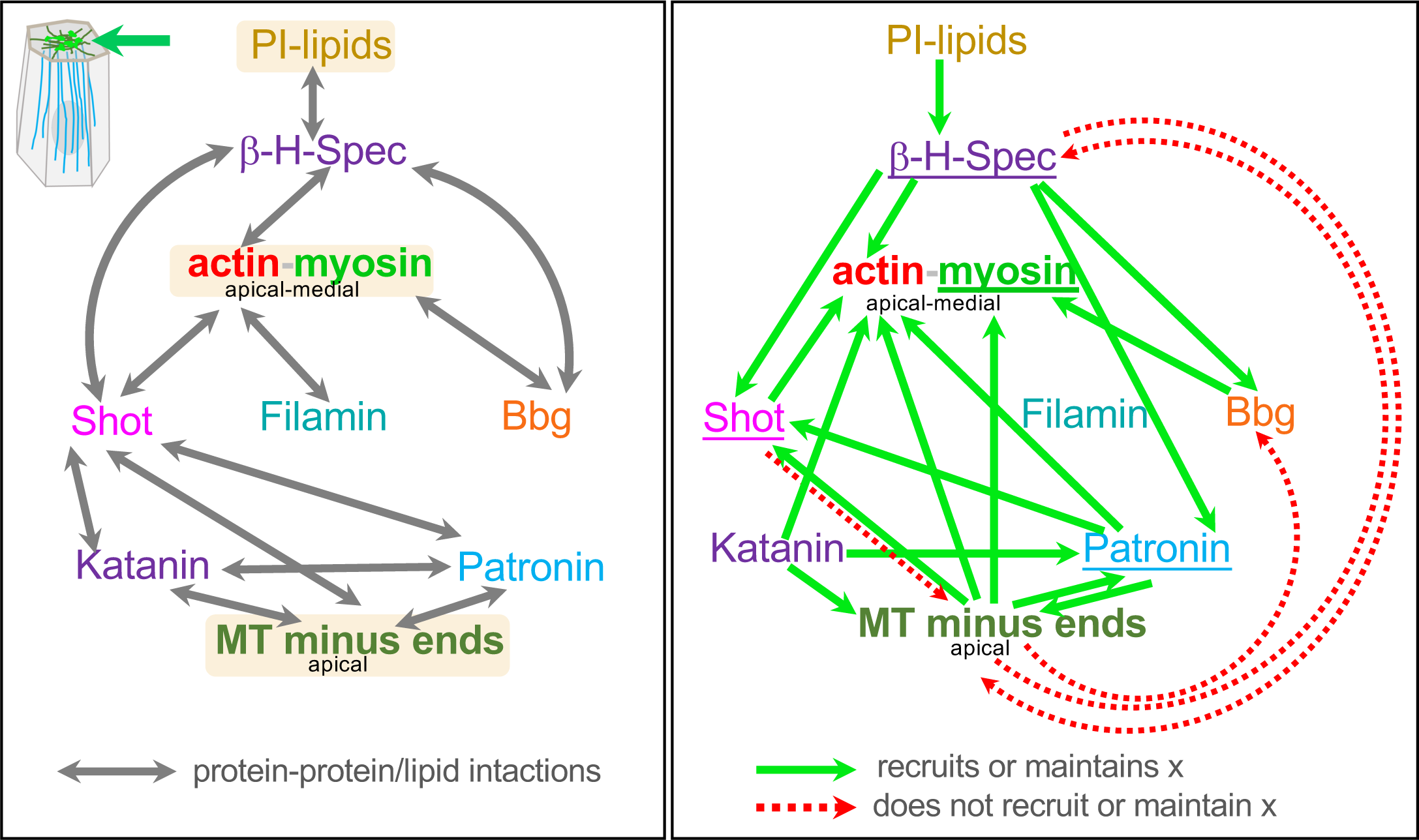
The apical-medial hub of cytoskeletal interactors. Schematics of the components identified that localise at the interface of apical-medial actomyosin and microtubule minus ends in placodal cells. Left panel illustrates protein-protein and protein-lipid interactions identified by us and others. Right panel shows where a component is required for other hub components’ recruitment or maintenance, with green arrows indicating that a component recruits another, and red dotted arrows indicating that a components is not required to recruit or maintain another.

Some components of the hub, we have shown previously, depend on the microtubule cytoskeleton, or rather the apical minus-ends of the longitudinal microtubule bundles, for their recruitment or maintenance, such as actomyosin, Patronin and Shot (Booth *et al*., 2014; Gillard *et al*., 2021). By contrast, β-H-Spectrin’s and Bbg’s apical-medial localisation does not depend on microtubules. For β-H-Spectrin, this localisation rather appears to rely on its C-terminal β-H segment 33. The β-H-33 segment has in fact previously been shown to exhibit autonomous membrane association in late stage embryonic salivary glands, in part mediated by the PH domain contained within it (Williams *et al*., 2004). PH domains are known to interact with phosphoinositides, including PI(4,5)P_2_ and PI(3,4,5)P_3_ (Lemmon, 2007). Phosphoinositides have been originally described as displaying a clear polarised distribution in epithelial cells, with PI(4,5)P_2_ being associated with the apical membrane (Martin-Belmonte *et al*., 2007) and PI(3,4,5)P_3_ localising to and regulating the formation of the basolateral membrane (Gassama-Diagne *et al*, 2006). However, recent findings suggest roles for both PI(4,5)P_2_ and PI(3,4,5)P_3_ within the apical domain of epithelial cells in tissues undergoing morphogenesis in *Drosophila* (Claret *et al*., 2014; Miao *et al*., 2021; Pinal *et al*., 2006). We find that fluorescent probes reporting on the presence of both PI(4,5)P_2_ and PI(3,4,5)P_3_ label the apical-medial domain of constricting salivary gland placodal cells, suggesting that β-H-Spectrin might be recruited to the domain via interactions through the PH domain. Furthermore, the fact that overexpression of the full β-H-33 fragment competes with endogenous β-H-Spectrin localisation at the apical-medial hub, leading to major morphogenetic defects, while overexpression of the β-H-33 segment without its PH domain does not impair the apical-medial accumulation of endogenous β-H-Spectrin, also strongly supports this notion. We propose that Bbg is then recruited through its direct interaction with β-H-Spectrin (Forest *et al*., 2018). Thus, the hub overall seems to depend on recruitment signals and synergy from two directions, the plasma membrane as well as the minus ends of longitudinal microtubule bundles. Interestingly, even though Patronin and Shot are reduced in their apical-medial pool when βH-Spectrin is depleted, Patronin function remains enough to support the formation of the longitudinal array, illustrating that in the wild-type, both the longitudinal array as well as βH-Spectrin and its interactors are key to apical constriction.

A further role suggested for β-H-Spectrin might also suggest a further functionality of the hub as a whole: β-H-Spectrin has been shown to be involved in mechanotransduction, by regulating the Hippo pathway (Fletcher *et al*, 2015) or endothelial mechano-responses (Mylvaganam *et al*, 2022). With pulsatile contractile cycles of actomyosin activity during apical constriction clearly exerting forces within a cell and via junctional contacts to neighbouring cells, β-H-Spectrin might also participate in mechanical responses of cells within the placode that help coordinate cell behaviours. Thus, we suspect that many important roles for the spectrin cytoskeleton remain to be uncovered, with our analysis of β-H-Spectrin function within the apical-medial hub illustrating one dynamic role.

## Acknowledgments

The authors would like to thank the following people; for reagents and fly stocks: Claire Thomas, Debbie Andrew, Daniel St. Johnston, Magali Suzanne, Alexandre Djiane, Markus Affolter; for use of the otracks software: Guy Blanchard.

This work was supported by the Medical Research Council, as part of United Kingdom Research and Innovation (also known as UK Research and Innovation) [MRC file reference number U105178780]. For the purpose of open access, the MRC Laboratory of Molecular Biology has applied a CC BY public copyright licence to any Author Accepted Manuscript version arising’.

## Author contribution

Conceptualisation, K.R. and G.G..; Methodology, K.R., G.G.; Investigation, K.R., G.G; Writing-Original Draft, K.R., G.G.; Funding Acquisition, K.R.

## Declaration of Interests

The authors declare no competing interests.

---

**Supplemental Figure S1.**
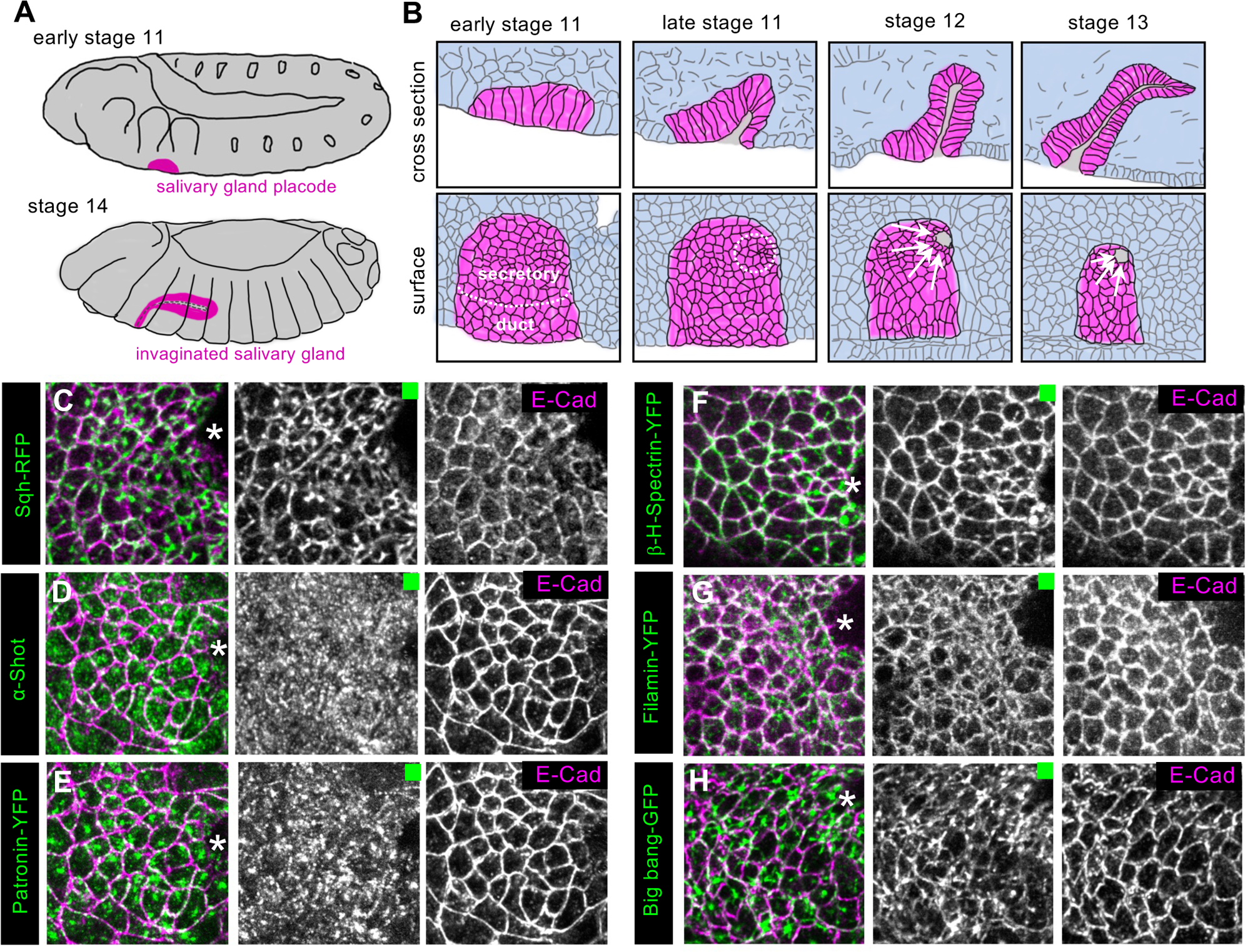
An apical-medial hub of proteins during apical constriction in tubulogenesis. **A** Schematic of *Drosophila* embryos at early stage 11 and stage 14 illustrating the position of the salivary gland placode at early stage 11 (magenta) and invaginated salivary gland at stage 14 (magenta), respectively. **B** Cross section and surface view schematics of salivary gland placodal cells (magenta) during the invagination process. Secretory cells begin to constrict in the dorsal posterior corner of the placode (dotted white circle) with cells continuously moving towards the pit through cell intercalations and to then apically constrict and invaginate (white arrows). **C-H** Individual channel panels corresponding to the panels shown in Figure 1 D-I. **C** Sqh-RFP, **D** anti-Shot, **E** Patronin-YFP, **F** β-H-Spectrin-YFP, **G** Filamin-YFP, **H** Bbg-GFP, all in green. Cell outlines are labelled for E-Cadherin (magenta). Asterisks indicate the position of the invagination point. Note that **D** and **E** correspond to a triple-labeling for Shot, Patronin-YFP and E-Cadherin.

**Supplemental Figure S2.**
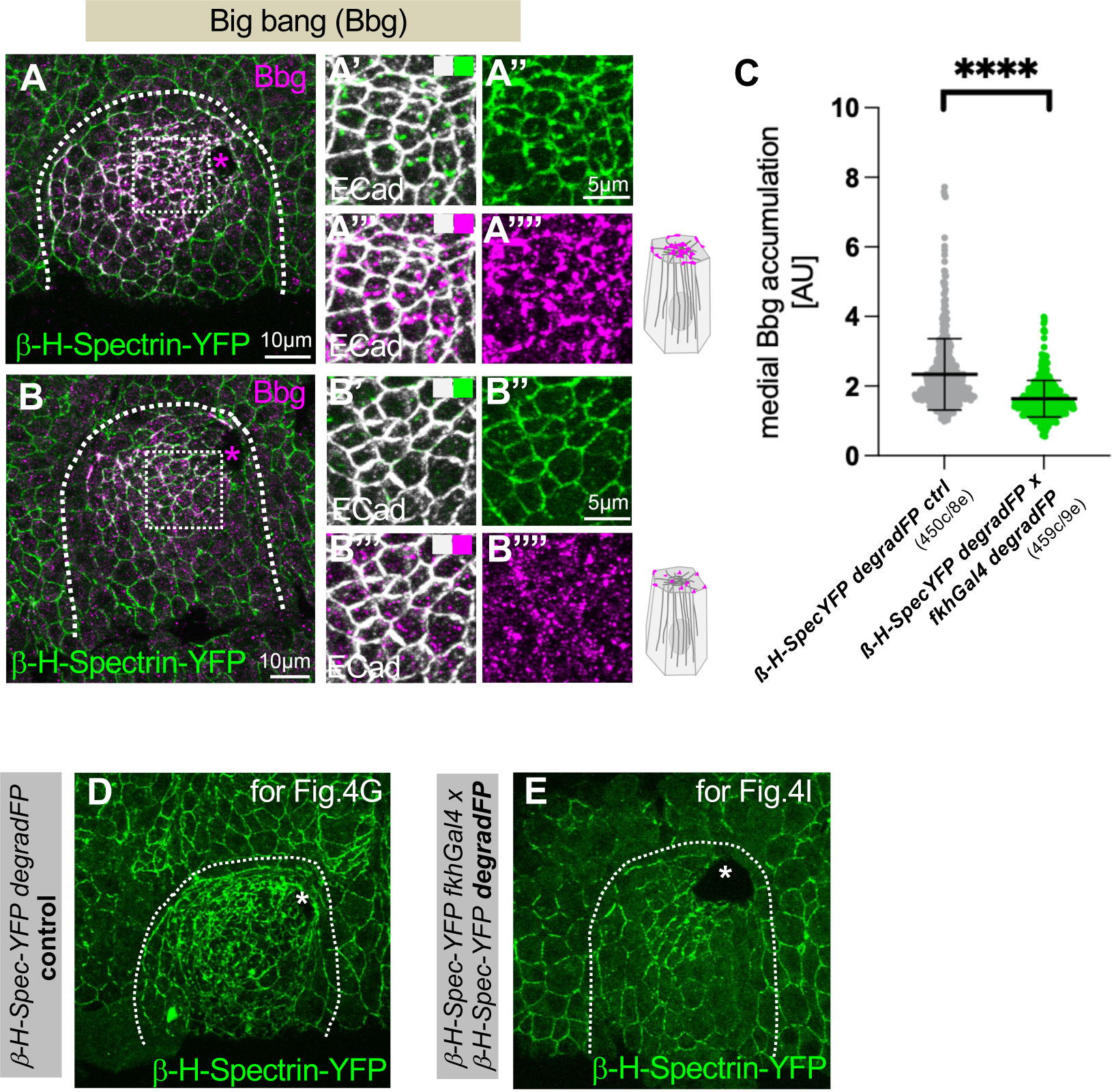
Loss of β-H-Spectrin leads to loss of the apical-medial hub. **A-C** β-H-Spectrin-YFP degradation (**B-B’’’’**) leads to a reduction in apical-medial Bbg foci (anti-Bbg, magenta) compared to control (**A-A’’’’**). β-H-Spectrin-YFP is in green and E-Cadherin to label apical cell outlines is in white. **A’-A’’’’** and **B’-B’’’’** are higher magnifications of the white boxes marked in **A** and **B**, respectively. **C** Quantification of apical-medial Bbg in placodal cells in control (*β-H-Spectrin-YFP fkhGal4 control;* 450 cells from 8 embryos) and β-H-Spectrin depleted (*β-H-Spectrin-YFP fkhGal4* x *β-H-Spectrin-YFP degradFP;* 459 cells from 9 embryos) placodes. Shown are mean +/−SD, statistical significance was determined by two-sided unpaired Mann-Whitney test as p<0.0001. **D,E** β-H-Spectrin-YFP in control ((*β-H-Spectrin-YFP fkhGal4;***D**) and when degraded ((*β-H-Spectrin-YFP fkhGal4* x *β-H-Spectrin-YFP degradFP*; **K**) for the BBg analysis shown in Fig.4 G-J. Asterisks indicate the position of the invagination pit; dashed lines mark the boundary of the salivary gland placode.

**Supplemental Figure S3.**
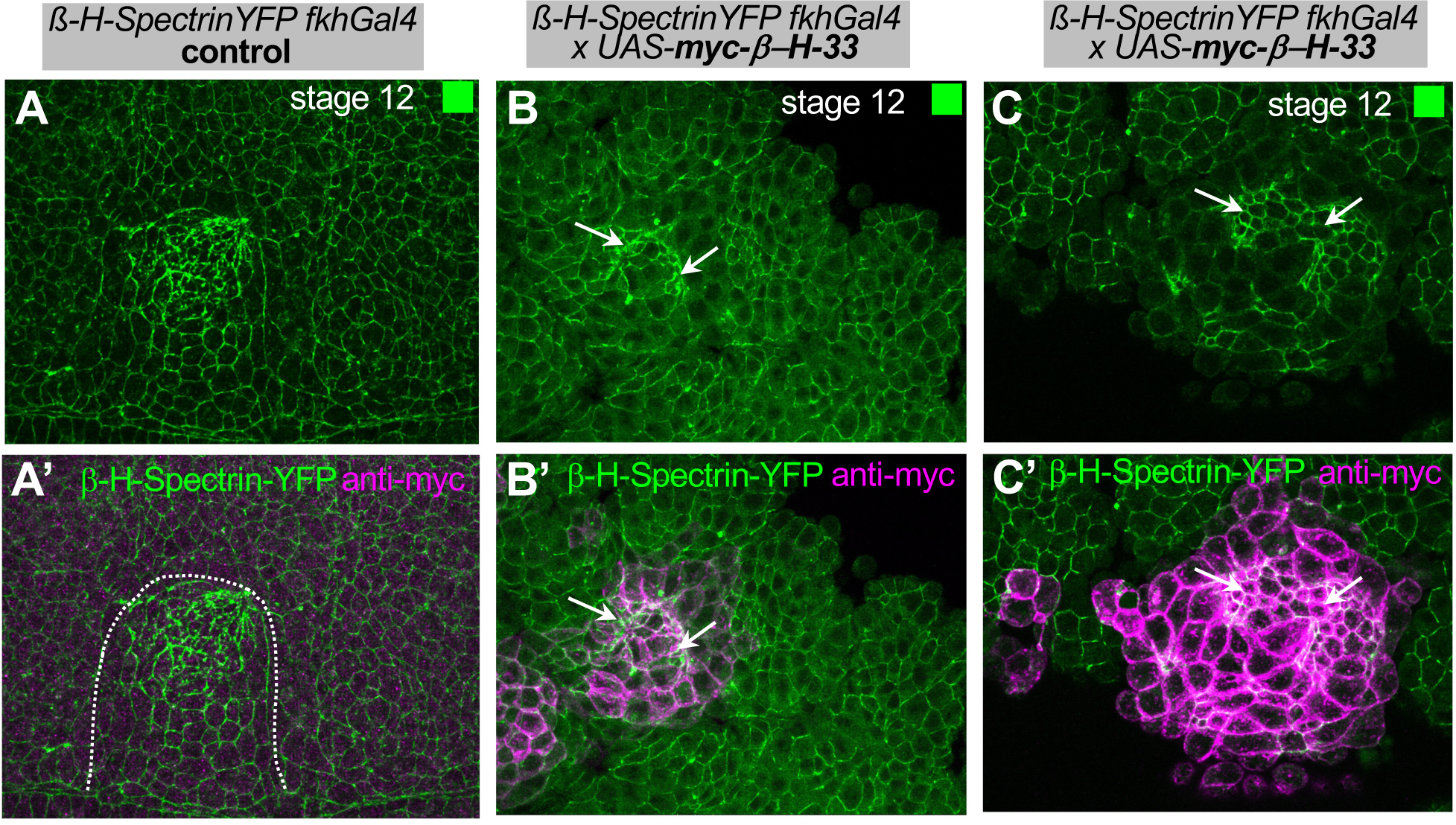
β-H-Spectrin’s localization to and function in the apical-medial hub depends on its PH domain. **A-C’** Overexpression of *UAS-myc-β-H-33* using *fkhGal4* at embryonic stage 12 in comparison to control as shown in Figure 6 H, I, J. Shown here are the β-H-Spectrin channel (**A-C**, green) and the overlay of β-H-Spectrin (**A’-C’**, green) and anti-myc staining (**A’-C’**, mangenta) to reveal the overexpressed fragment. Arrows point to the remaining constricting cells when βH33 is overexpressed, the dotted line in **A’** marks the boundary of the placode, in **B’** and **C’** the anti-myc labelling indicates placodal cells due to the use of *fkhGal4*.

## Methods

### Drosophila stocks and genetics

The following fly stocks were used in this study: from Bloomington Stock Centre: *Daughterless-Gal4* (*Da-Gal4*; #27608); Bbg-GFP (y[1] w[*]; ; Mi{PT-GFSTF.1}bbg[MI02662-GFSTF.1]/TM6C, Sb[1] Tb[1] #60187). Furthermore *β-H-SPectrin-YFP*/Kst-YFP (w1118; PBac{681.P.FSVS-1}kstCPTI002266) [Kyoto Stock Centre]; Patronin::TagRFPattp40[22H02-C] (Takeda *et al*, 2018); *sqh-TagRFPt[9B]* (Ambrosini *et al*., 2019); *fkh-Gal4* on chromosome III (Henderson & Andrew, 2000); *Patronin-YFP (w1118; Patronin-YFP/Cyo*) (Nashchekin *et al*., 2016); *UAS-deGradFP (w; If/Cyo; UAS>NSlmb-vhhGFP4/TM6b*) (Caussinus *et al*., 2012); *UAS-Spastin* on X (Sherwood *et al*, 2004); P{w+mc kstβH33.Scer\UAS.T:Hsap\MYC=pβH33} and P{w+mc kstβH33ΔPH.Scer\UAS.T:Hsap\MYC=pβH33ΔPH} (Williams *et al*., 2004). (Booth *et al*., 2014; Gillard *et al*., 2021)

The following strains of transgene combinations were generated in the context of this study:

*Patronin-RFP; β-H-SPectrin-YFP*
*Sqh-RFP; ;β-H-SPectrin-YFP*
*EGFP-Shot;β-H-SPectrin-YFP*
*β-H-SPectrin-YFP fkh-Gal4*
*UAS-Spastin; ;β-H-SPectrin-YFP*
*Pat-RFP;β-H-SPectrin-YFP fkh-Gal4*
*UAS-degradFP;β-H-SPectrin-YFP*
Genotypes analysed are indicated in the figure panels and legends.

### Embryo Immunofluorescence Labelling, Confocal, and Live Analysis

Embryos were collected on apple juice-juice plates and processed for immunofluorescence using standard procedures. Briefly, embryos were dechorionated in 50% bleach, fixed in 4% MeOH-free formaldehyde, and devitellinised in a 50% mix of 90% EtOH and Heptane or MeOH and Heptane. They were then stained with phalloidin or primary and secondary antibodies in PBT (PBS plus 0.5% bovine serum albumin and 0.3% Triton X-100). anti-E-Cadherin (DCAD2, 1:10 dilution) and anti-CrebA (CrebA Rbt-PC, 1:1000) antibodies were obtained from the Developmental Studies Hybridoma Bank at the University of Iowa; anti acetylated α-tubulin (clone 6-11B-1, 1:500; Sigma); anti-Shot (1:1000; (Röper & Brown, 2003)). The following secondary antibodies were used at 1:200: Alexa Fluor® 488 AffiniPure Donkey Anti-Rabbit IgG (H+L) (711-545-152); Cy™3 AffiniPure Donkey Anti-Rabbit IgG (H+L) (711-165-152); Alexa Fluor® 647 AffiniPure Donkey Anti-Rabbit IgG (H+L) (711-605-152); Cy™3 AffiniPure Donkey Anti-Mouse IgG (H+L) (715-165-151); Cy™5 AffiniPure Donkey Anti-Mouse IgG (H+L) (715-175-151); Cy™3 AffiniPure Goat Anti-Rat IgG (H+L) (112-165-167); Alexa Fluor® 647 AffiniPure Donkey Anti-Rat IgG (H+L) (712-605-153); Cy™5 AffiniPure Donkey Anti-Guinea Pig IgG (H+L) (706-175-148) were from Jackson ImmunoResearch Laboratories. Donkey anti-Rabbit IgG (H+L) Highly Cross-Adsorbed Secondary Antibody, Alexa Fluor Plus 405 (A48258); Goat anti-Mouse IgG (H+L) Highly Cross-Adsorbed Secondary Antibody, Alexa Fluor 350 (A-21049); Donkey anti-Mouse IgG (H+L) Highly Cross-Adsorbed Secondary Antibody, Alexa Fluor Plus 488 (A32766); Goat anti-Rat IgG (H+L) Cross-Adsorbed Secondary Antibody, Alexa Fluor 488 (A-11006) were from Invitrogen, and rhodamine-phalloidin (1:500) was from Thermofisher (R415). Samples were embedded in Vectashield (Vectorlabs H-1000).

Images of fixed samples were acquired on an Olympus FluoView 1200 (with the FV10-ASW v04.02 software) or a Leica SP8 inverted microscope (LAS X software) equipped with 405nm laser line for four-colour imaging as z-stacks to cover the whole apical surface of cells in the placode. Z-stack projections and orthogonal sections were assembled in ImageJ or Imaris (Bitplane), 3D rendering was performed in Imaris.

For live time-lapse imaging, the embryos were dechorionated in 50% bleach, rinsed in water and attached to a coverslip with the ventral side up using heptane glue and covered with Halocarbon Oil 27. Time-lapse sequences of embryos of the genotypes *Sqh-RFP;;β-H-YFP Patronin-RFP; ;β-H-YFP* were acquired every 6.1s and 17.4s, respectively, on a spinning disc set-up as z-stacks, while embryos of the genotype *EGFP-Shot; β-H-SPectrin-YFP* were imaged every 7s on a Zeiss 880 inverted microscope (Zen 2.3 SP1 FP3 v14.0.20.201 software) with a 40x/1.3NA Oil objective as a single confocal slice, using linear unmixing to allow imaging of both fluorophores.

Z-stack projections to generate movies were assembled in ImageJ or Imaris.

### Quantifications

#### Cell segmentation and apical area analysis

For the analysis of apical cell area, images of fixed embryos of late stage 11/early stage 12 placodes, labelled with DE-Cadherin to highlight cell membranes and with dCrebA to mark salivary gland fate, were analysed. Cells were segmented in confocal image stacks, with cell analysis software (otracks, custom software written in IDL, from L3 Harris Geospatial, https://www.l3harrisgeospatial.com/Software-Technology/IDL; code availabe on request from Dr Guy Blanchard) as used and published previously (Blanchard *et al*, 2009; Blanchard *et al*, 2010; Booth *et al*., 2014; Butler *et al*, 2009). Briefly, the shape of the curved placode surface was identified in each z-stack as a contiguous ‘blanket’ spread over the cortical signal. Quasi-2D images for cell segmentation containing clear cell cortices were extracted as a maximum intensity projection of the 1 or 1.5 µm thick layer of tissue below the blanket. These images were segmented using an adaptive watershed algorithm. Manual correction was used to perfect cell outlines for fixed embryos. Only cells of the salivary placode were used in subsequent analyses and were distinguished based on dCrebA staining.

#### Medial accumulation of phalloidin, Bbg, β-H-SPectrin-YFP, Patronin-RFP and Shot

Images were taken of salivary gland placodes and surrounding tissue at late stage 11/early stage 12. Maximum intensity projections of the apical surface of placodal cells were generated using 3-5 optical section separated by 1 µm each in z. For each embryo analysed, fluorescence measurements were made for all secretory cells within the placode except cells close to the actomyosin cable. The medial and junctional values were measured after drawing the cells outlines (7-pixel wide line) with a home-made plugin in Fiji, available on request. 10 cells were similarly analysed in the surrounding tissue.

For Kst-YFP, Shot and phalloidin quantifications, the graphs display the medial accumulation corresponding to the ratio between the medial intensity for each cell in the placode divided by the mean intensity of the 10 cells outside the placode. For Patronin-RFP, due to the noisy labeling in the surrounding epidermis, the graph displays the ratio between Patronin-RFP medial staining versus junctional staining for each cell in the placode.

### Statistics and Reproducibility

Significance was determined using two-tailed Student’s *t*-test, non-parametric Mann-Whitney test for non-Gaussian distribution, unpaired t-test with Welch’s correction for data with unequal standard deviations, and Kolmogorov-Smirnov (K-S) test for the comparison of cumulative distributions.

Tests used are indicated in Figure legends.

## Supplemental Movie Legends

**Supplemental Movie 1. Dynamic behaviour of myosin and β-H-Spectrin in placodal cells.**

Time lapse movie of a salivary gland placode of an embryo with the genotype *Sqh-RFP;; β-H-Spectrin-YFP*, with Sqh-RFP in magenta and β-H-Spectrin-YFP in green. Time interval is indicated.

**Supplemental Movie 2. Dynamic behaviour of Shot and β-H-Spectrin in placodal cells.**

Time lapse movie of a salivary gland placode of an embryo with the genotype *ShotEGFP; β-H-Spectrin-YFP*, with ShotEGFP in magenta and β-H-Spectrin-YFP in green. Time interval is indicated.

**Supplemental Movie 3. Dynamic behaviour of Patronin and β-H-Spectrin in placodal cells.**

Time lapse movie of a salivary gland placode of an embryo with the genotype *PatroninRFP; β-H-Spectrin-YFP*, with PatroninRFP in magenta and β-H-Spectrin-YFP in green. Time interval is indicated.

## Notes

### Competing Interest Statement

The authors have declared no competing interest.

